# SNP profile for quantitative trait nucleotide in populations with small effective size and its impact on mapping and genomic predictions

**DOI:** 10.1101/2023.02.16.528829

**Authors:** Ignacy Misztal, Ivan Pocrnic, Daniela Lourenco

**Affiliations:** Department of Animal and Dairy Science, University of Georgia, Athens, GA 30602, USA; The Roslin Institute, The University of Edinburgh, EH25 9RG, Edinburgh, UK

**Keywords:** distribution of SNP effects, sequence data, effective population size, single-step GBLUP

## Abstract

In animal populations, increasing the SNP density by incorporating sequence information only marginally increases prediction accuracies. To find out why, we used statistical models and simulations to investigate the profile or distribution of SNP around Quantitative Trait Nucleotides (QTN) in populations with small effective population size (N_e_). A QTN profile created by averaging SNP solutions around each QTN was similar to the shape of expected pairwise linkage disequilibrium (PLD) based on N_e_ and genetic distance between SNP, with a distinct peak for the QTN. Populations with smaller N_e_ showed lower but wider QTN profiles; however, adding more genotyped individuals with phenotypes dragged the profile closer to the QTN; the QTN profile was higher and narrower for populations with larger compared to smaller N_e_. Assuming the PLD curve for the QTN profile, 80% of the additive genetic variance explained by each QTN is contained in 8 “Stam” segments (one segment = 1/4N_e_ Morgans), corresponding to 1.6 Mb in cattle, and 5 Mb in pigs and broiler chickens. With such large segments, identifying QTN is difficult even if all of them are in the data and the assumed genetic architecture is simplistic. Additional complexity in QTN detection arises from confounding of QTN profiles with signals due to relationships, overlapping profiles with closely-spaced QTN, and spurious signals due to imputation errors. However, small N_e_ allows for accurate prediction with large data even without QTN identification because QTN are accounted for by QTN profiles if SNP density is sufficient to saturate the segments.

## INTRODUCTION

Sequence data brings the opportunity to search for and use causative variants (quantitative trait nucleotides; QTN) for genomic predictions in animal and plant breeding or the prediction of human polygenic risk scores. If most of the QTN were known, predictions would be more accurate and persistent. If those QTN were similar across populations (or breeds) and had similar substitution effects, an accurate admixed evaluation would be possible. So far, the use of putative QTN in genomic predictions has sometimes resulted in slightly increased accuracy but not always − e.g., see a review by Hayes and Daetwyler (2019). An increase of up to 5% in accuracy was reported when eliminating large SNP around putative QTN (VanRaden *et al*. 2017).

Potential QTN or SNP close to QTN used for genomic predictions are typically identified by genome-wide association studies (GWAS). A standard tool for traditional GWAS is a model where one marker is analyzed at a time as a fixed effect, whereas the polygenic effect is accounted for by fitting a relationship matrix among individuals under mixed models (Kennedy *et al*. 1992). Such a model is also known as efficient mixed-model association expedited or EMMAX (Kang *et al*. 2010). Fitting a relationship matrix based on pedigree or genotypes reduces spurious signals due to population structure because it assumes individuals may share a considerable proportion of genes (Kennedy 1991; Kang *et al*. 2010). Alternatively, many recent studies are adopting Bayesian regression methods like BayesB (Meuwissen *et al*. 2001) or BayesR (Erbe *et al*. 2012) that consider all SNP jointly as random and estimate the effect of a SNP conditionally to all the other SNP. The SNP with larger signals are considered markers of nearby QTN. While the single SNP models use p-values to determine SNP significance, the joint SNP models usually estimate fractions of explained variance per segment of the genome, e.g., 1Mb, and may use power and false discovery rates as arbitrary approaches to compare methods. While the golden standard for putative QTN is ≥ 1% of explained additive genetic variance (e.g., Chen *et al*. 2017), the origin of a 1 Mb segment is not clear. Another GWAS method gaining momentum in animal and plant populations is the single-step GWAS or ssGWAS (Wang *et al*. 2012; Aguilar *et al*. 2019). This method estimates all SNP simultaneously, provides variance explained by each SNP together with a significance test (i.e., p-values equivalent to EMMAX; Gualdron-Duarte et al., 2014; Aguilar et al., 2019) and is based on single-step GBLUP (ssGBLUP; Aguilar *et al*. 2010; Christensen and Lund 2010), which allows using information on all individuals concurrently, independently of the pedigree, phenotyping, and genotyping status. Single-step GBLUP is the standard tool for genomic predictions in farm animal populations (Tsuruta *et al*. 2011; Christensen *et al*. 2014; Lourenco *et al*. 2015) and was recently applied to the UK Biobank data (Truong *et al*. 2020).

Independently of the method, an essential question for identifying QTN in animal and plant populations is how the smaller effective population sizes limit the resolution of GWAS compared to humans. The genome comprises blocks or chromosome segments inherited from founders, separated by junctions (Fisher 1949; Fisher 1954); those junctions are swapping spots for the founder origin of segments. For randomly mating populations of constant size, the number of junctions is a function of effective population size (Ne) and genome length (L) (Stam 1980). Changing population size (i.e., population growth, division, or bottleneck) strongly affects the number of junctions in small but not large populations (Chapman and Thompson 2002). Junctions define the genome segments; therefore, the inheritance is by segments, not by individual genes or QTN. A limited number of segments has important implications in GWAS as the segment size affects the resolution (Berisa and Pickrell 2016). For instance, Wang *et al*. (2012) found that the correlation between the effects of QTN and one adjacent SNP was lower than between a segment of 16 adjacent SNP and QTN. Assuming a genome size of 3 Gb and the number of junctions of 10,000 in animals or 1 million in humans, the segment size and, subsequently, resolution of GWAS would be approximately 3 kb in humans and 300 kb in animals. GWAS results do provide evidence of this limited resolution. Manhattan plots on individual SNP are usually noisy, and a common strategy is to smooth out noise by combining variance explained by SNP segments of, say, 1 Mb (Funkhouser *et al*. 2020). A limited number of segments constrains the dimensionality of genomic information for populations with small Ne and allows for high prediction accuracy of the genomic merit even with fewer data (Pocrnic *et al*. 2019).

While the above studies suggest that Ne limits the resolution of GWAS, it is possible to envisage a scenario where identifying QTN is feasible despite limited Ne. Assume a small number of QTN, all present in the data, a large number of phenotypic records, and an additive QTN model. If only QTN were used in SNP-BLUP or GBLUP-based models, the prediction accuracy would be close to 100%, as found by Fragomeni *et al*. (2017) or PÉrez-Enciso *et al*. (2015). Then, QTN could be determined by an exhaustive search for a set of SNP that results in the best predictive power, which means almost 100% predictive accuracy. This scenario is hypothetical as most traits are likely to be controlled by many QTN, with most below the detection threshold.

In this paper, we study the pattern of SNP distribution around a QTN in populations with varying Ne and numbers of genotyped individuals with phenotypes. This pattern is referred to as the QTN profile and helps understand the resolution of GWAS and its impact on genomic selection. Investigating the QTN profile around a QTN in GWAS and determining whether it is a function of effective population size likely requires large datasets and many replicates. In this study, we determine the profile of QTN using data simulated to minimize the sampling variance. We also discuss the implications of this profile in methods used for genomic predictions.

## MATERIALS AND METHODS

### Chromosome segments and pairwise linkage disequilibrium

The expected number of chromosome segments given by Stam (1980) is 4 NeL, where Ne is the effective population size, and L is the genome length in Morgans. Subsequently, the average size of a Stam segment (s) is 1/4 Ne Morgan. Additionally, the expected pairwise linkage disequilibrium (PLD), represented by r^2^, was defined by Sved (1971) as

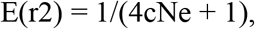

where c is the genetic distance between two SNP in Morgans. The plot of r^2^ as a function of c in terms of Stam segments is given in Figure 1. The interval where E(r^2^) declines to 66%, 50%, 33%, and 20% corresponds to 1, 2, 4, and 8 Stam segments, respectively, as shown in the Appendix. For more discussions on LD and chromosome segments, see Goddard and Meuwissen (2005).

**Figure 1.**
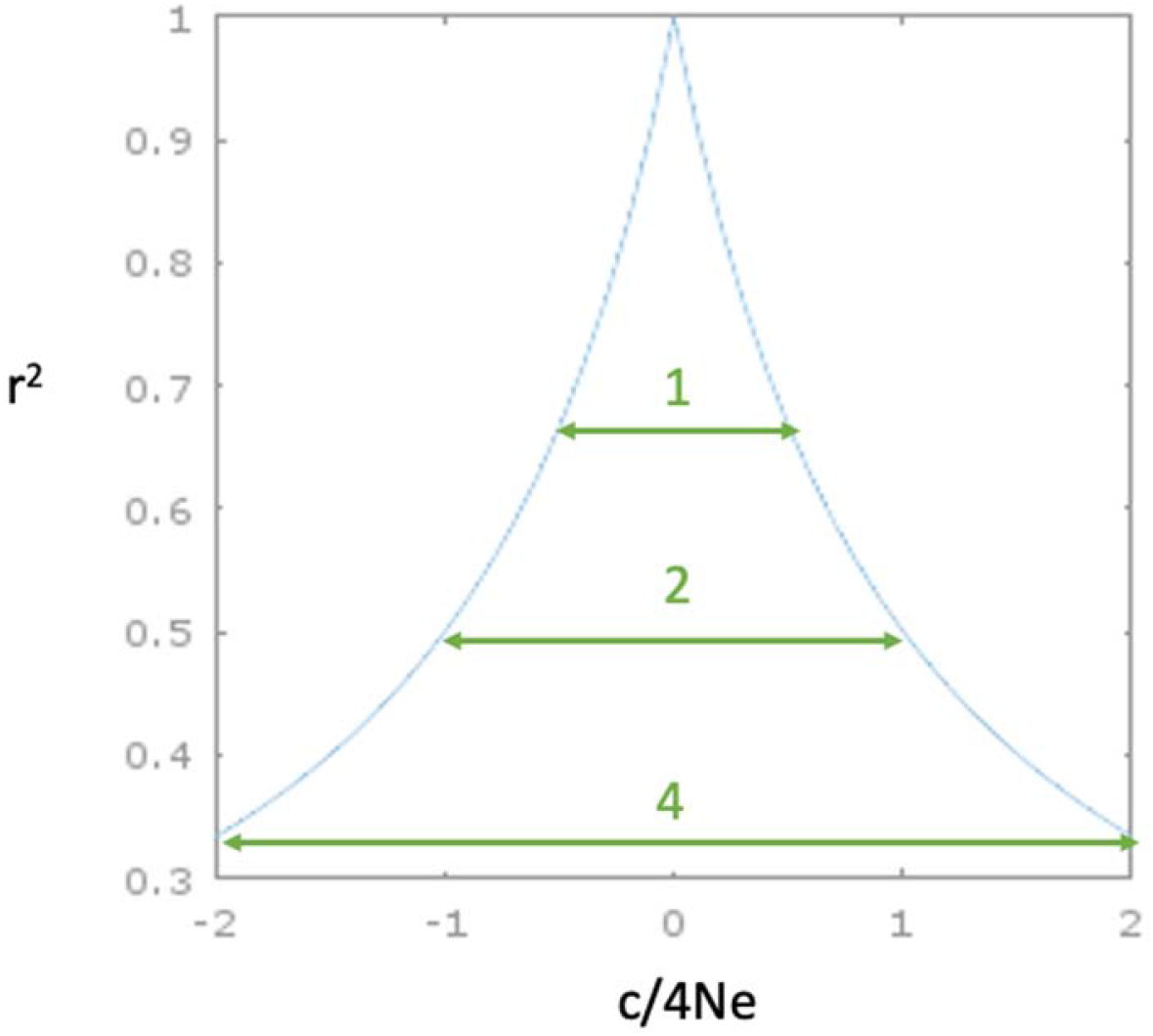
Expected value of pairwise linkage disequilibrium (r^2^) as a function of distance from the QTN in Stam segments.

### Phenotypic and genotypic data

Data for this study were simulated using the AlphaSimR package (Faux *et al*. 2016; Gaynor *et al*. 2019) and was run using the R version 3.4.4 (R Core Team). A historical population was generated using the default options of the coalescence simulator as implemented in the package, except that the final effective population size (Ne) was fixed to either 60 or 600. The first recent generation (base population) was created using 6000 individuals from the historical population, followed by nine generations of random mating to ensure that the Ne remained as close as possible to that simulated in the historical population. The number of mating males and females per generation was set to 15 and 1000 for Ne=60, 175 and 1000 for Ne=600, and 15 and 3000 for Ne=60, but with three times more individuals per generation. Further on, these scenarios will be referenced as NE60, NE600, and NE60_3x, respectively.

In all scenarios, the simulated genome had 10 chromosomes with equal lengths of 100 cM each. To limit the computations, 50,000 biallelic SNP markers were used, equally spaced along the chromosomes, resulting in 50 SNP per cM. Equal placement of genetic markers was achieved by modifying the default genetic map in AlphaSimR. Each chromosome harbored 10 QTN that were assigned the same additive effect and placed in the same locations across the 10 chromosomes, corresponding to the locations of actual SNP markers. The QTN were separated by at least 500 SNP (≈10cM) to reduce interference. Two progenies of equal sex ratio were created per mating, resulting in either 2,000 (NE60 and NE600) or 6,000 (NE60_3x) individuals per generation. Phenotypes for a quantitative trait were generated assuming a heritability of 0.5 and with a single record per individual. Pedigree and phenotypes were recorded for all 24,000 (NE60 and NE600) or 60,000 (NE60_3x) animals in the recent populations. Genomic information was available for the last three generations, i.e., 6,000 (NE60 and NE600) or 18,000 (NE60_3x) genotyped individuals.

### Single-step genome-wide association analysis

Most of the GWAS methods assume that genotypes and phenotypes are available in the same individuals; however, in livestock populations, those two sources of information may be available in different individuals. Because of that, we used single-step genome-wide association (ssGWAS) (Wang *et al*. 2012; Aguilar *et al*. 2019) under the following model:

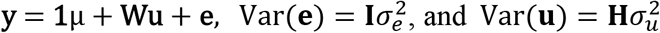

where **y** is a vector of phenotypes, μ is an overall mean, **u** is a vector of random additive animal effects, **e** is a vector of random residuals, **W** is an incidence matrix relating observations in **y** to additive genetic effects in **u**, **H** is a realized relationship matrix, 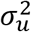 is the additive variance, and 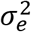 is the residual variance. Variance components were assumed to be known using the base population simulation parameters. The realized relationship matrix **H** combines pedigree and genomic relationships, with the inverse as in Aguilar *et al*. (2010):

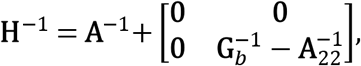

where **A** is the numerator relationship matrix for all animals included in the analysis, and **A**_22_ is the pedigree relationship matrix for genotyped animals. To ensure the matrix was invertible, the initial **G** was blended prior to inversion as **G**_*b*_ **=** *α***G** + (1-*α*)**A**_22_, with *α* = 0.95, and the initial **G** defined as in VanRaden (2008):

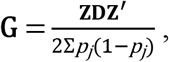

where ***Z*** is a matrix of allele content centered for allele frequencies, *p_j_* is the allele frequency for marker *j* in the current genotyped population, and **D** is a diagonal matrix of weights for SNP markers. All SNP were assumed to have equal weight; therefore, **D** was an identity matrix (**I**).

After computing genomic estimated breeding values (GEBV), SNP effects (a) were obtained as in Wang *et al*. (2012):

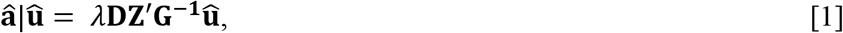

where *λ* is a ratio of SNP to additive variance used in the data simulation. The SNP effects used in this study were scaled as 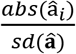. P-values for SNP were computed based on Aguilar *et al*. (2019):

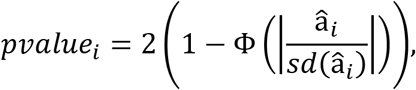

with Φ being the cumulative standard normal function and 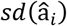 the square root of prediction error variance (PEV) of the *i^th^* SNP effect. Prediction error variance for each SNP effect was:

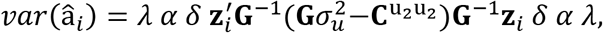

where *δ* = (1 − *ρ*/2), and *ρ* is the average difference between all elements of **G** and **A**_22_, which is known as the tuning parameter used to adjust the genetic base of **G** to **A**_22_ (Vitezica *et al*. 2011). All SNP that passed the Bonferroni threshold of 10^−6^ were considered statistically associated with the simulated trait. Computations were by the BLUPF90 software suite (Misztal *et al*. 2014).

### Pooled SNP effects for QTN profiling

Because QTN and SNP were simulated at the same positions across the 10 chromosomes, we averaged the effects of 100 SNP closest to the causative variants, i.e., 50 SNP upstream and downstream of the QTN, in the 10 chromosomes to assess the QTN profiles. This is equivalent to averaging segments of approximately 1 cM for a population with Ne equal to 60.

### Data availability

The authors state that all data necessary for confirming the conclusions presented in the article are represented fully within the article.

## RESULTS AND DISCUSSION

### Genome-wide Manhattan plots

Manhattan plots of standardized SNP effects for NE60, NE60_3x, and NE600 are in Fig. 2. Because 100 equidistant QTN were simulated with identical effects, there was an expectation of observing roughly 100 similar peaks. However, only a few peaks could be identified, and the plots had low resolution (i.e., noisy plots). Visually, the number of signals above the noisy region increased from NE60 to NE600, with NE60_3x in between the two. Some differences in signals from QTN could be due to changes in gene frequencies because of natural selection or drift; no directional selection was simulated and the mating was random.

**Figure 2.**
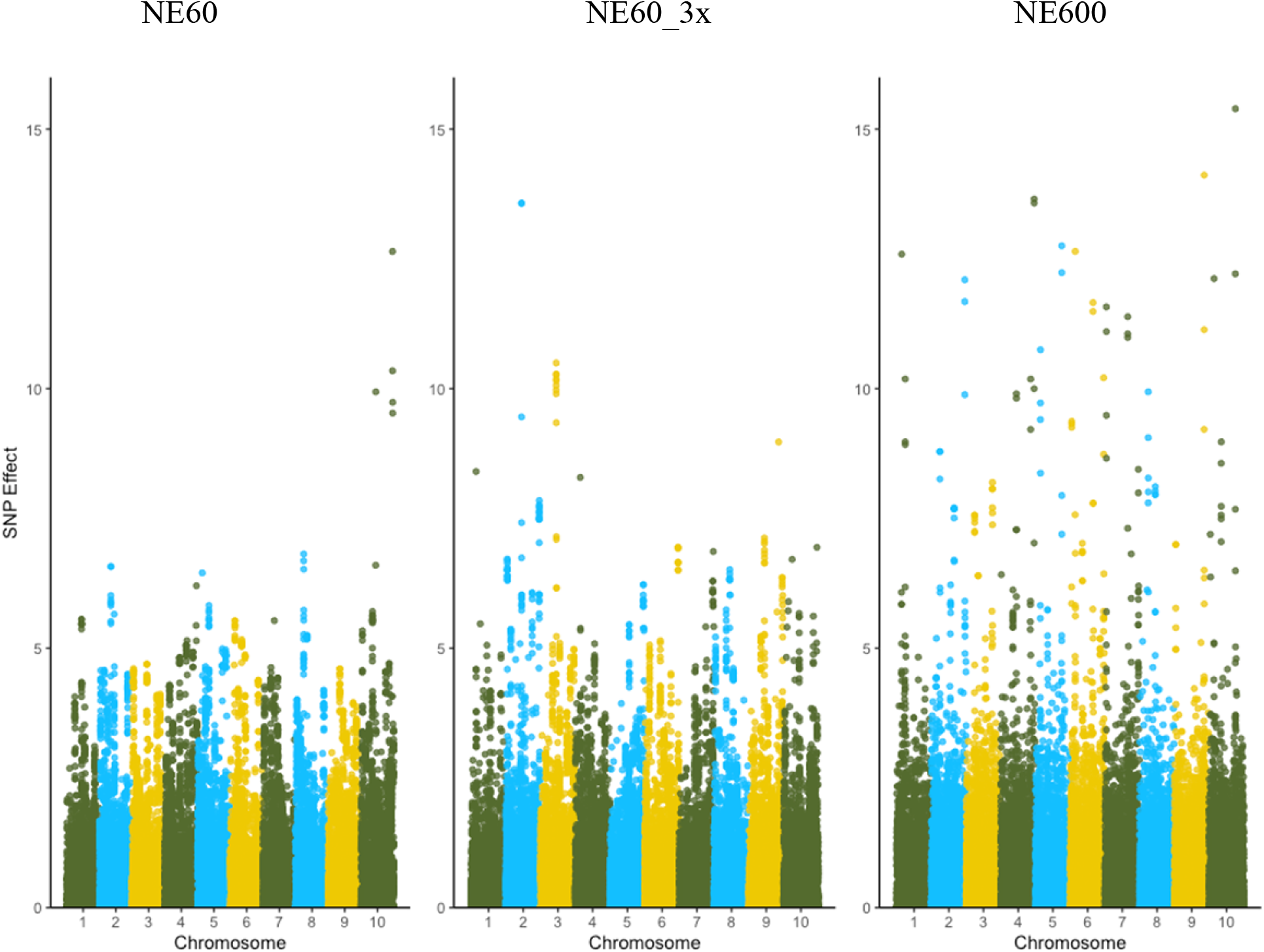
Manhattan plots for SNP standardized effects computed for datasets with effective population size 60 (NE60), with the same effective population size but 3 times more data (NE60_3x), and with effective population size 600 (NE600).

To assign significance to the signals while accounting for population structure, we recreated Manhattan plots for p-values using the scale of −log 10 (*p-value*), which are shown in Fig. 3. The number of significant SNP, i.e., the SNP above the threshold of 6 on the −log 10 scale, was the smallest with NE60, larger with NE60_3x but with similar background noise, and the largest for NE600, with more negligible background noise. Figures 2 and 3 were comparable, indicating that the plots based on SNP effects and p-values are visually similar. While Fig. 2 is based on the value of the SNP effect, Fig. 3 is based on a function of the SNP effect adjusted for its standard deviation (SD). The plots are approximately proportional when the SDs are similar, although not linearly. More significant SNP were captured with a larger sample size or larger Ne. The association signals are less noisy with more individuals and, subsequently, with more recombination.

**Figure 3.**
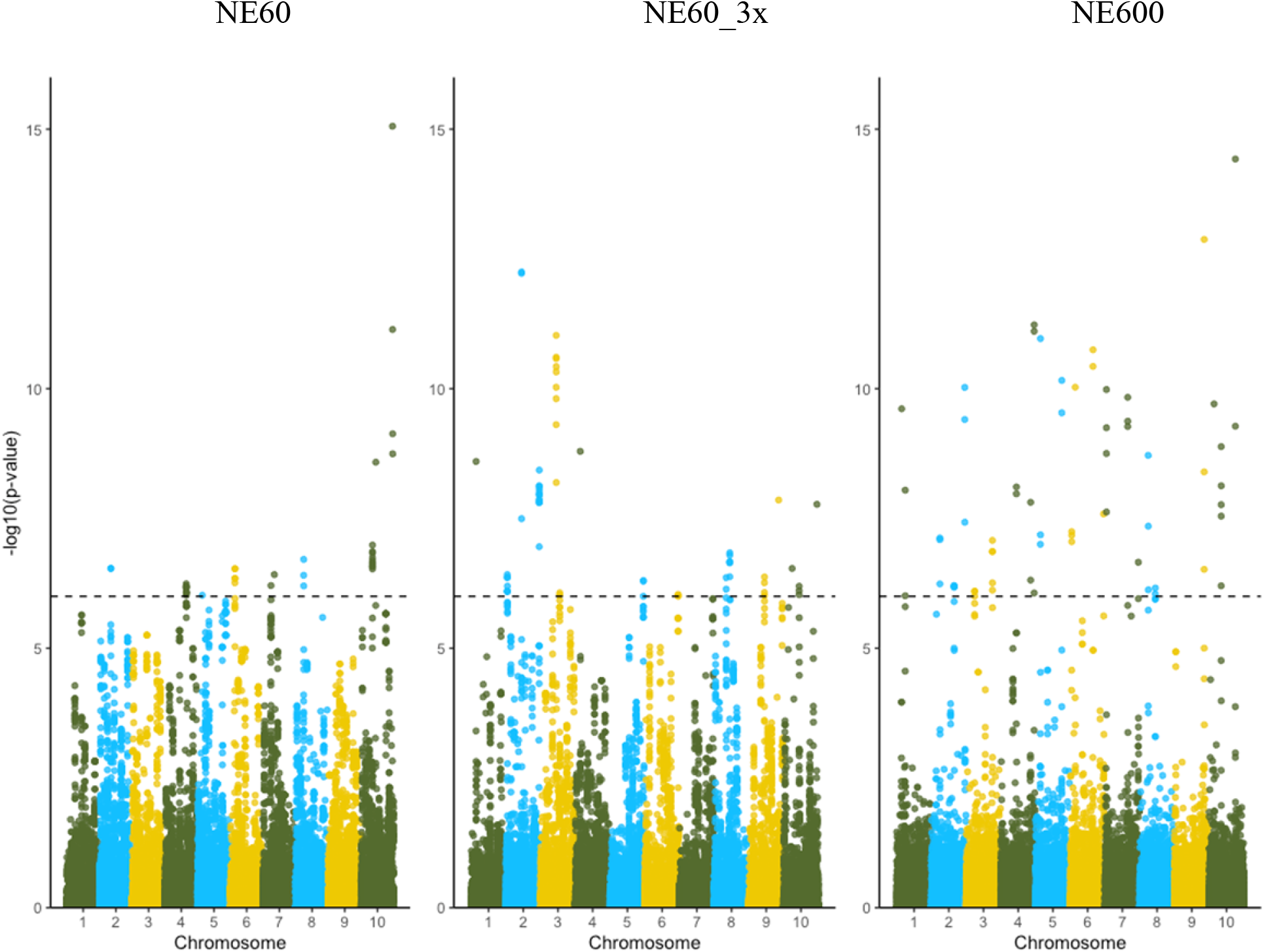
Manhattan plots for p-values computed for datasets with effective population size 60 (NE60), with the same effective population size but 3 times more data (NE60_3x), and with effective population size 600 (NE600).

### Detection of QTN in a single chromosome

To visualize signals close to QTN, Fig. 4 shows Manhattan plots of SNP effects from the first chromosome with locations of each of the 10 QTN identified. Again, 10 signals of approximately the same value were expected. However, all plots were noisy, and only a few of the 10 simulated QTN (dotted lines) had a trail of SNP, although with a smaller magnitude than the simulated effects. Signals increased when the data size or Ne increased. Because of the high noise level, QTN profiles were not evident from these plots.

**Figure 4.**
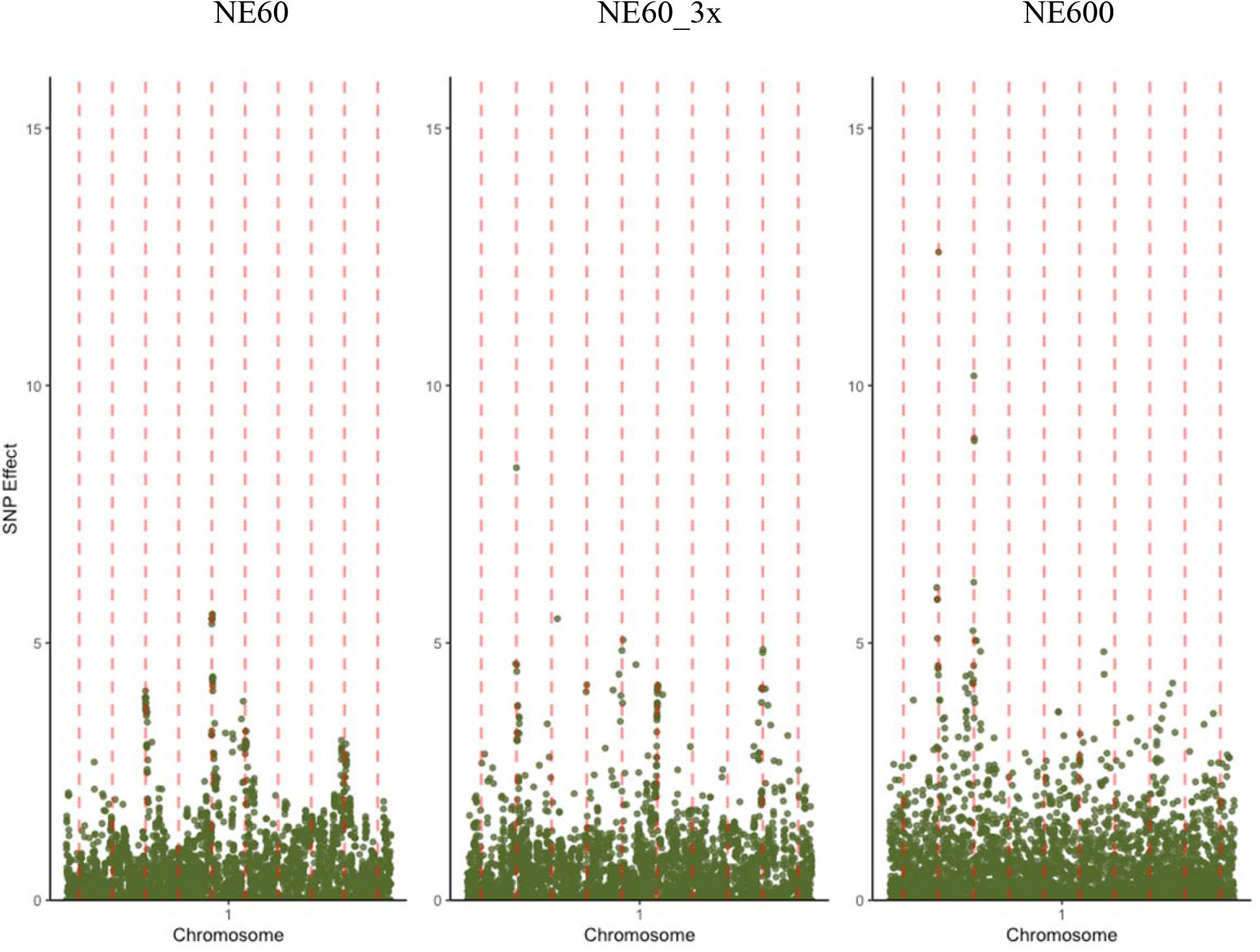
Manhattan plots for SNP effects - first chromosome only - computed for datasets with effective population size 60 (NE60), with the same effective population size but 3 times more data (NE60_3x), and with effective population size 600 (NE600). Vertical red lines indicate locations of causative SNP.

### Pooled SNP effects

As all QTN were simulated with the same effect, it was possible to reduce the noise by averaging the effects of 50 SNP upstream and downstream from QTN, equivalent to averaging segments of approximately 1 cM for Ne equal to 60, as shown in Fig. 5. In all scenarios, the maximum peak response was at the true QTN position and had values from about 3.0 to 3.5 SD, with the remaining SNP showing a distribution with a sharp peak, similar to a Laplace distribution as in Bayesian models (De los Campos *et al*. 2009). For NE60 and NE60_3x, the averaged response had a similar distribution, although the variability around the curve was higher for NE60, and the peak was higher for NE60_3x. For NE600, the average profile was lower and narrower, with the peak at the QTN.

**Figure 5.**
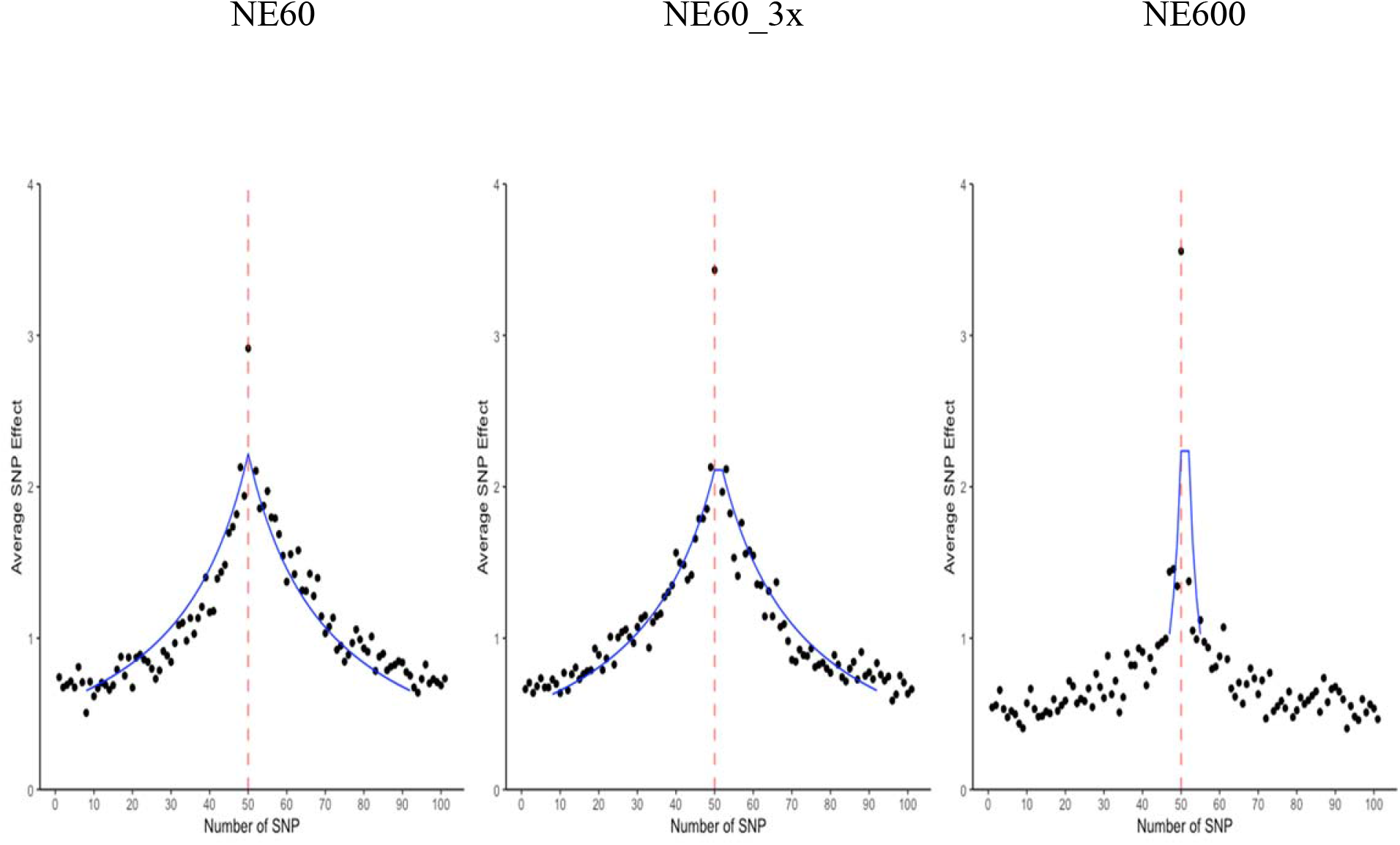
Profile of causative SNP or distribution of SNP around causative SNP, in standard deviation units, computed for datasets with effective population size 60 (NE60), with the same effective population size but 3 times more data (NE60_3x), and with effective population size 600 (NE600). Vertical red lines indicate locations of simulated QTN. Blue curves indicate the best fit with the pairwise linkage distribution curve; R^2^ for the fit excluding the QTN is 0.89, 0.83 and 0.23 for NE60, NE60_3x, and NE600, respectively.

Above the background level of 1 SD of average SNP effect, the profile was about 60 SNP or 1 cM wide for NE60 and about 10 SNP or 0.2 cM wide for NE600. The five times wider profile with NE60 compared to NE600, despite the 10 times difference in Ne, could be due to ignoring the profile below 1 SD. Using the formula by Stam (1980), the number of independent chromosome segments (ICS), equivalent to the number of genome segments, is 4NeL or 2400 for NE60 and 24,000 for NE600. Assuming 50k SNP, this would correspond to a segment of approximately 20 SNP for NE60 and 2 SNP for NE600. A wider profile in GWAS than that of ICS means that the profile spans many ICS.

The true value of each QTN is about 22 SD of a single SNP, whereas the maximum for a single SNP is 15 (Fig. 2), with the average in Fig. 5 around 3.5. This means that the identified QTN explained, on average, only a small fraction of about (3.5/22)^2^ = 2.5% of each QTN standardized effect. Based on the pooled graphs (Fig. 5), the QTN profile elicits a wide response, with the width of the response being a function of the effective population size. Additionally, the highest response is for the actual QTN, if present in the data, and it is higher with more data. Shrinkage reduces the effect of QTN; the shrinkage would be stronger with more SNP as the variance of an individual SNP is inversely proportional to the number of SNP.

### Pairwise linkage disequilibrium

We hypothesize that the QTN profile is a function of pairwise linkage disequilibrium (PLD; r2) as defined by Sved (1971): E(r2) = 1/(4cNe + 1), where c is the genetic distance between two SNP, expressed in Morgans. Such a formula is visualized in Fig. 1, with numbers represented as the length of one segment as derived from the formula by Stam (1980), where one segment is 1/4Ne Morgans. The expected PLD decays to 67% for an interval of a single Stam segment, 50% for an interval of two Stam segments, 33% for an interval of 4 Stam segments, and 10% for 19 Stam segments. In this study, one Stam segment would be about 20 SNP for Ne equal to 60 and 2 SNP for Ne equal to 600. Subsequently, the PLD would decay to 33% for an interval of 80 SNP with NE60 and NE60_3x, and 8 SNP with NE600. Assuming that PLD is the real QTN profile, SNP in 2 (4, 8) Stam segments would correspond to 50% (66%, 80%) of the total response to one QTN (see Appendix).

### Fitting pairwise linkage disequilibrium

Figure 5 shows Manhattan plots for the SNP effects with the superimposed PLD curves, displaying similar shapes. For intervals of ± 2 Stam segments around the QTN (80 SNP for NE60 and NE60×3, and 8 SNP for NE600) and excluding the QTN, the fit was precise for NE60 (R^2^ = 0.89) and NE60_3x (R^2^ = 0.83), and less so for NE600 (R^2^ = 0.23). A slightly poorer fit with NE60×3 compared to NE60 could be due to less shrinkage of the QTN effect. Less fit with NE600 is due to insufficient crossovers to saturate an 8 SNP interval. With 3k animals and a genome length of 10 Morgans, there are only 30k crossovers, or approximately one every 2 SNP, insufficient for a good fit. Therefore, larger data and more SNP would be required to improve the fit with NE600.

### Components of Manhattan plot

The predictive ability of GBLUP-based methods is mainly independent of the number of QTN (Lourenco *et al*. 2017; Takeda *et al*. 2021) and is attributed to exploiting differences between the expected and realized relationships (Vanraden 2008). SNPBLUP, and indirectly GBLUP, partially account for QTN as shown in this study; however, the signals due to QTN are affected by shrinkage and noise, the latter partly due to estimation error, genotyping errors, and a small number of SNP. Assuming that PLD is a good predictor of QTN profiles, it is possible to identify components of the Manhattan plot, as shown in Fig. 6–8. The plots are composed of signals due to relationships, LD with QTN following the PLD curve, actual QTN if present in the data, and signals due to noise because of estimation error, a finite number of SNP, and a limited number of samples.

**Figure 6.**
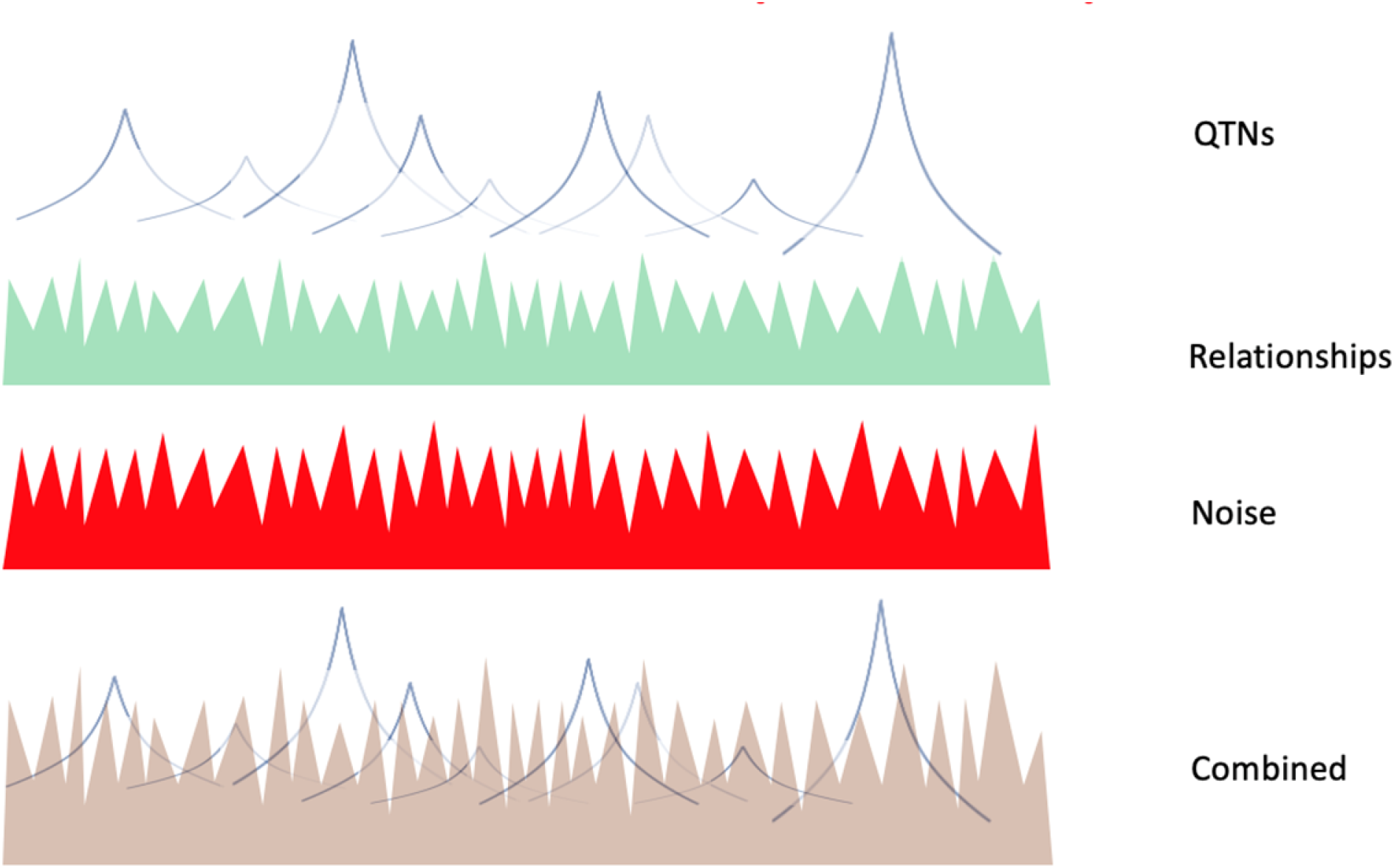
Components of a Manhattan plot and their composite plot for a small Ne and medium dataset.

While accuracies of genomic relationships are high with a typical number of SNP (SD < 0.5% with 40k SNP as in VanRaden 2008), signals due to relationships appear as semi-random noise as defined by formula [1] when all QTN are small. Signals due to QTN and QTN profiles are visible only when they are large enough to rise above the signals due to relationships and noise. Signals due to LD with QTN are wide for populations with small Ne, with 4 Stam segments accounting for up to 66% of QTN variance and 8 Stam segments accounting for up to 80% variance (see Appendix). The fraction of the QTN variance explained by the segments depends on Ne, the amount of data, and the distribution of the QTN effects. As only a fraction of QTN with similar effects were observed in this study, there is a strong confounding of QTN signals with other signals.

While Fig. 6 shows the Manhattan plot for a medium size dataset and small effective population size, Fig. 7 and 8 show the plots with a large dataset and many or few large QTN. With a small dataset, signals due to relationships prevail as signals due to QTN are small because of shrinkage, with a risk of pseudo-random peaks being interpreted as QTN or markers to QTN. In a study involving a fertility trait with low heritability in a small population with about 2000 dairy cattle (Kiser *et al*. 2019), the Manhattan plots lacked resolution, and many SNP were labeled as causative variants. With a large dataset, the estimation error and shrinking are smaller (Jang *et al*. 2022a). When many large QTN are present and now account for a large fraction of the additive variation, signals due to relationships decrease (Fig. 7). When few large QTN are present, and QTN profiles explain a small fraction of the additive variance, signals due to relationships remain the dominant part of the Manhattan plot. In a study involving a large population of 36,000 high-accuracy bulls (Jiang *et al*. 2019), the Manhattan plots were clear and showed many peaks with precise LD patterns.

**Figure 7.**
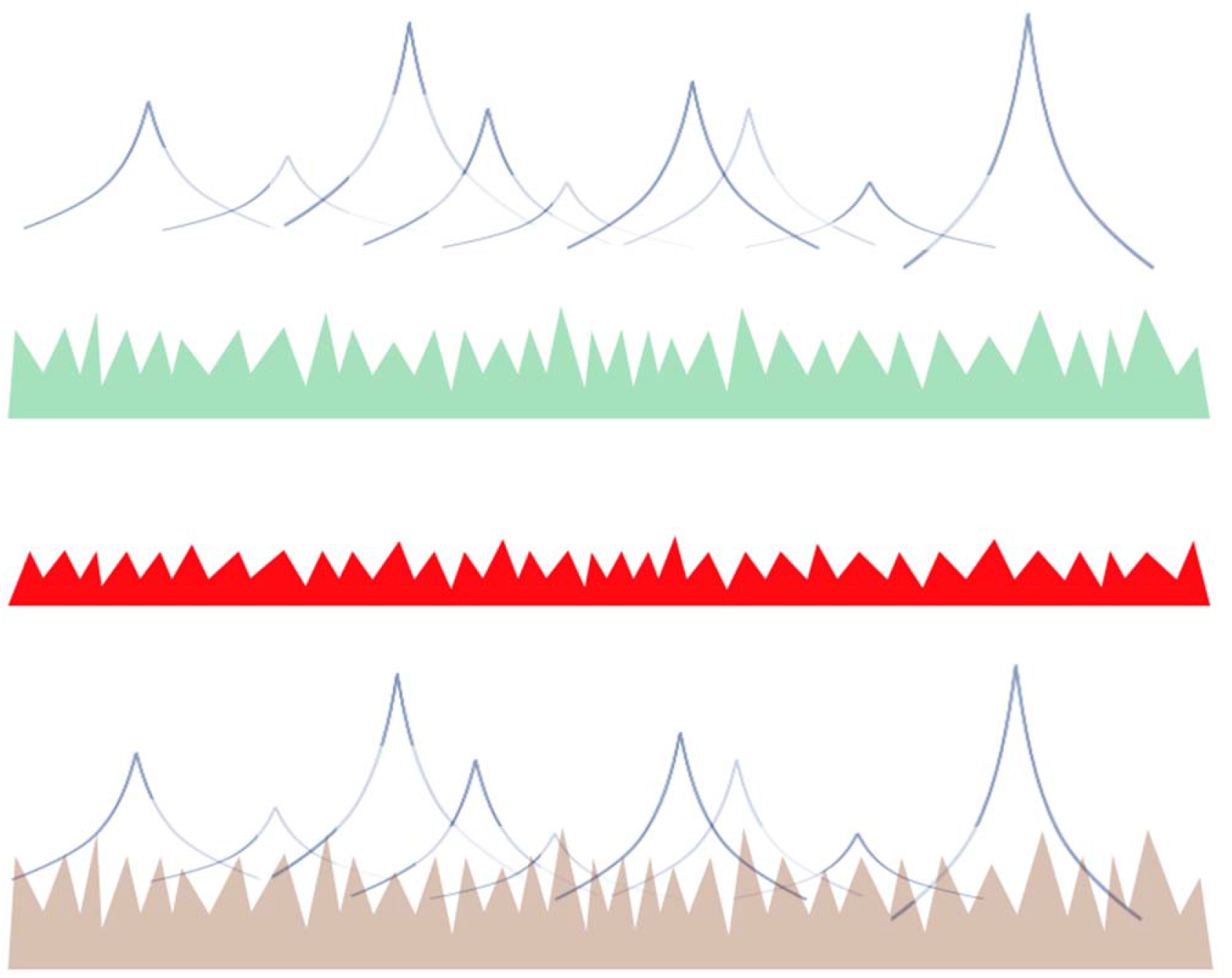
Components of a Manhattan plot and their composite plot for a small Ne and large dataset with many large QTL.

**Figure 8.**
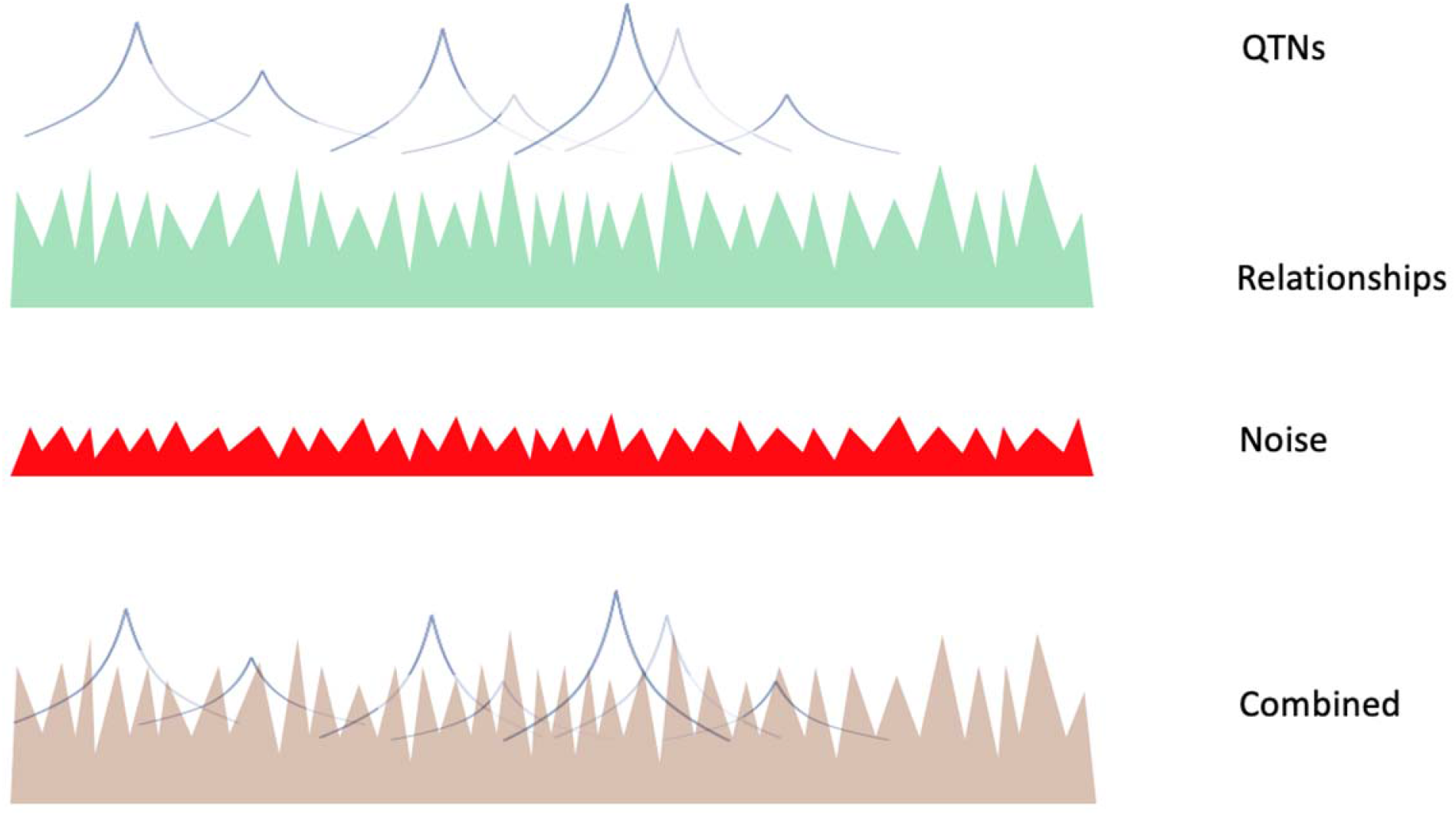
Components of a Manhattan plot and their composite plot for a small Ne and large dataset with few large QTL.

Figure 9 shows the Manhattan plot for a population with a very large effective size, e.g., humans. Signals due to relationships are very small, QTN profiles are very narrow, and the Manhattan plot is mainly composed of estimation error and very narrow profiles of SNP. Assuming genome length L=30 M, 8 Stam segments accounting for 80% of the QTN variance would be 2 Mb for a population with Ne=100 (e.g., cattle), 5 Mb for a population with Ne=40 (e.g., chicken and pigs), and only 20 kb for a population with Ne=10,000 (humans). When chosen experimentally by minimizing noise and maximizing information varied from 1 Mb in cattle (Buchanan *et al*. 2016) to 10 Mb in chicken (Stainton *et al*. 2017).

**Figure 9.**
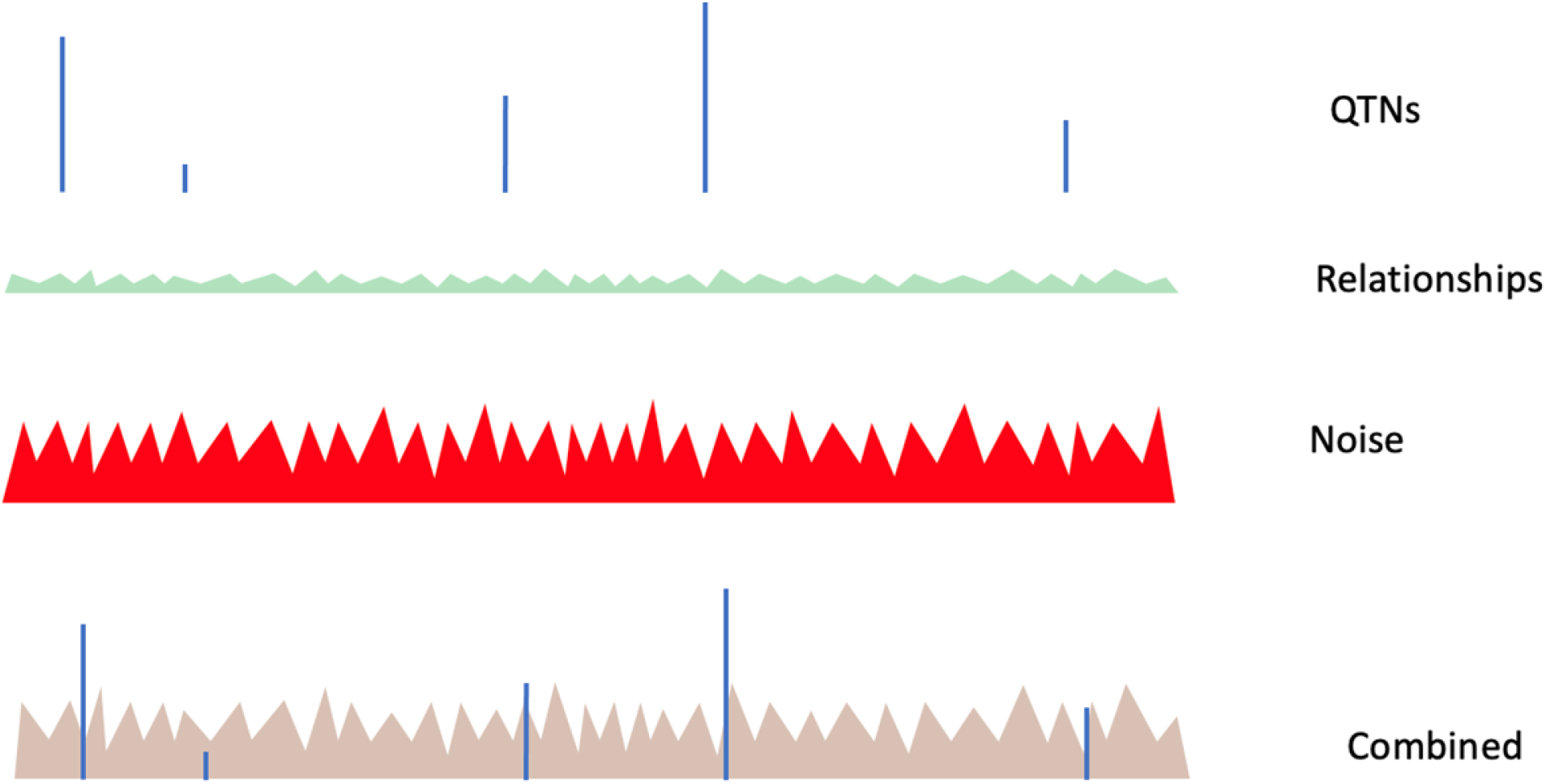
Components of a Manhattan plot and their composite plot for a large Ne.

With very large datasets, GBLUP or SNPBLUP incorporate large QTN by accounting for QTN profiles, as shown in Figure 10. Wang *et al*. (2012) looked at the prediction of QTN effects in a simulation study. The best estimates were not with the nearest SNP effect but with a sum of 16 nearby SNP, indicating the optimum window size of 16 SNP. With a small effective population size, QTN profiles of adjacent QTN are likely to overlap, and the observed peak in a Manhattan plot may be a composite of many QTN. The ability of GBLUP to account for QTN is a very valuable outcome for commercial genetic evaluations in plants and animals. Most models used for the genetic evaluation are multi-trait, and accounting for different QTN for each trait would lead to excessive computations (Tiezzi and Maltecca 2015).

**Figure 10.**
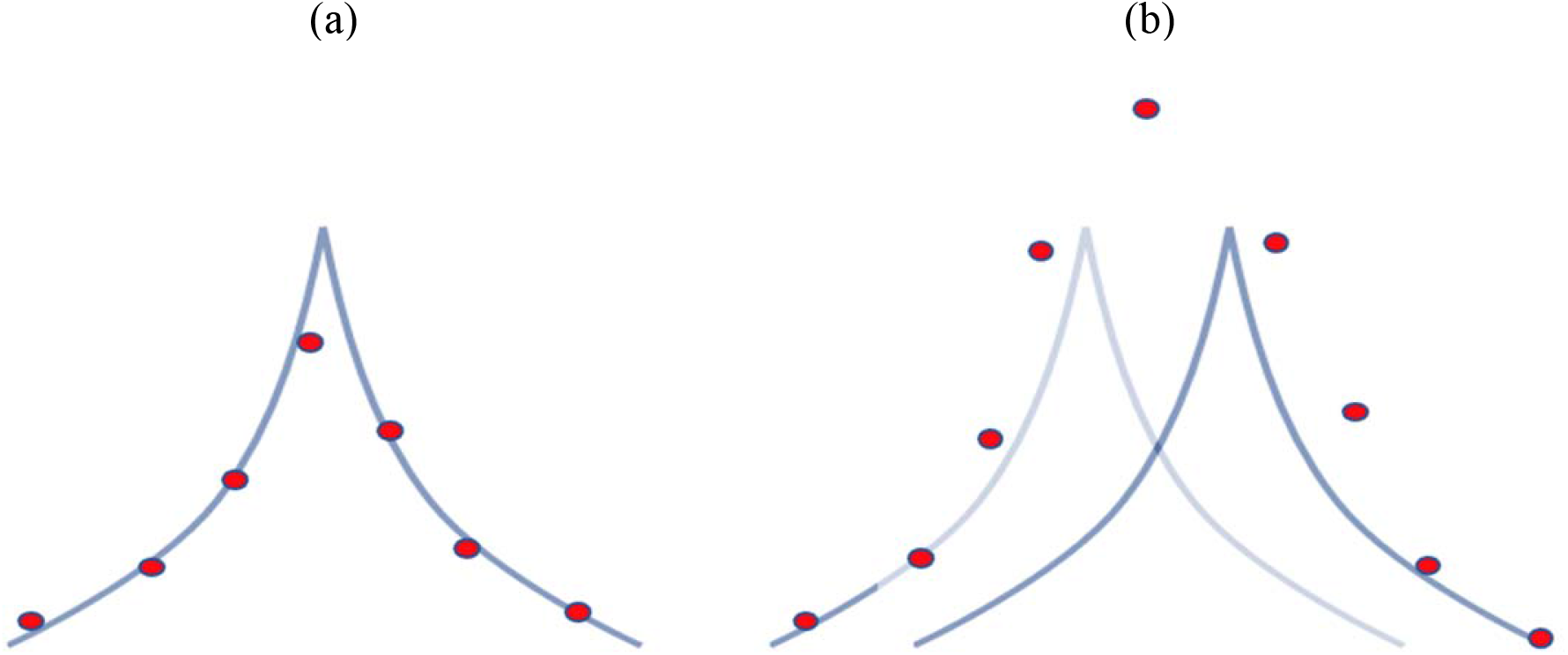
Accounting for QTN by GBLUP for single (a) and two close (b) QTN.

### Simulated versus real data

In our study, we used very strong assumptions to visualize the profile of QTN, including equal substitution effects for QTN, a small number of evenly spaced QTN, and equal recombination rate across the genome. In reality, most traits are complex and, therefore, controlled by a large number of not evenly spaced QTN, with only a few having a large effect size. Figure 11 shows hypothetical distributions of gene effects for selected and unselected populations. After intensive selection, most genes with a large effect become fixed, and large distributing genes are not fixed mainly because of pleiotropy, as documented by Georges *et al*. (2019). In the end, the only QTN of interest would be the top QTN not showing excessive pleiotropy. While this study looked at a single trait only, determining pleiotropy would require multi-trait GWAS.

**Figure 11.**
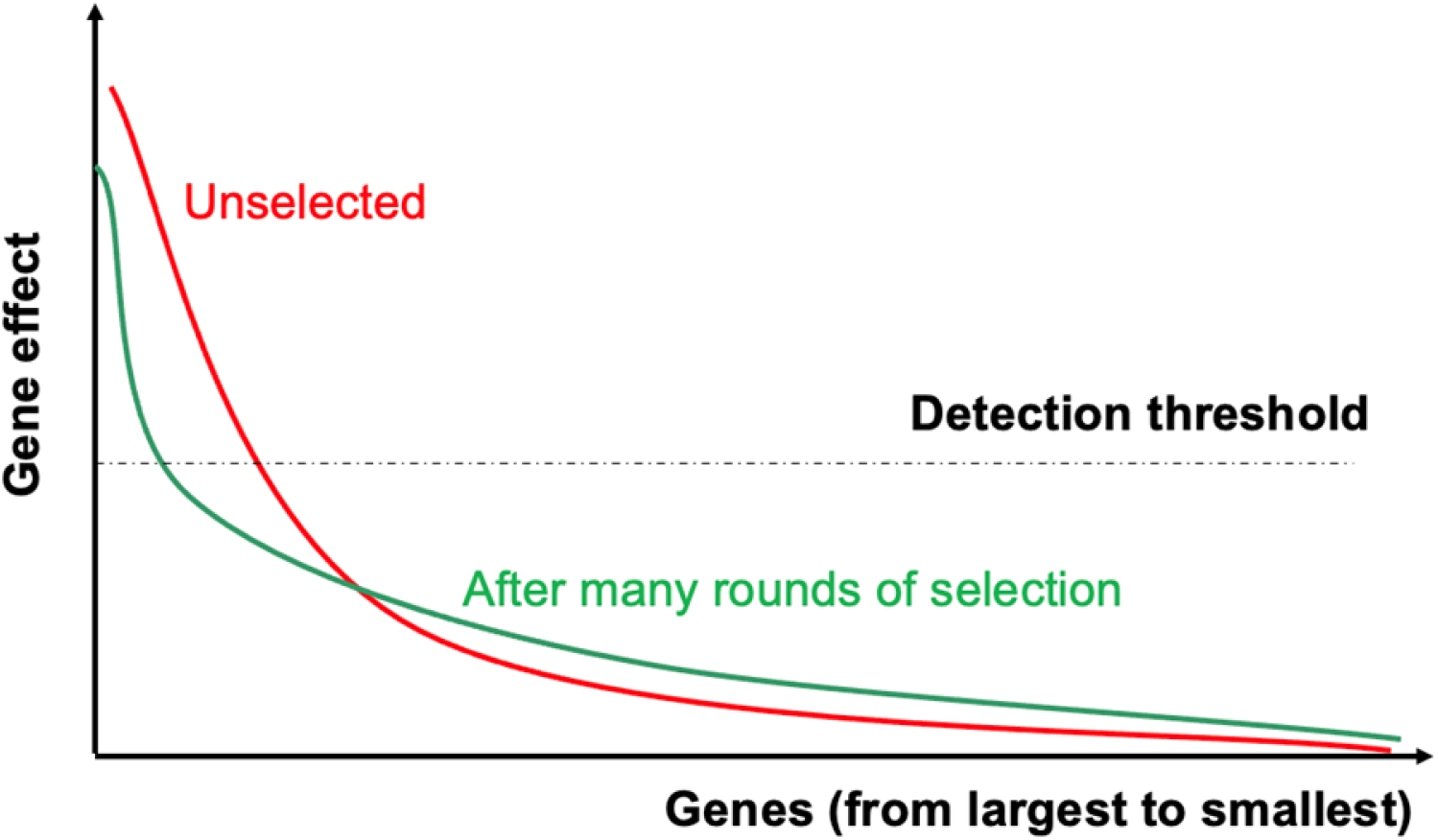
Distribution of gene effects for unselected traits and after many rounds of intensive selection.

### Implications for genomic selection

This study raises a question on the optimal SNP selection (or weighting of a genomic relationship matrix) based on statistical criteria applied by common methods, and its impact on accuracy on the genomic selection. Selected SNP can either be the actual QTN, markers to QTN as QTN profiles, markers due to relationship signals, or due to noise. The success of SNP selection also depends on the genetic architecture of traits (Zhang *et al*. 2016). With a few QTN, all of them can be identified and estimated well for high prediction accuracy. With a medium number of QTN, accurate predictions would require prior knowledge of the variance of QTN to avoid under or overprediction of the QTN effects. With many QTN, selected SNP would likely include only a few QTN (Fragomeni *et al*. 2017). BrØndum *et al*. (2015) stated that aside from knowing the variance of the QTN, knowing their positions helps to assign the variance to the correct variant, avoiding either shrinkage or inflation; shrinkage is less important with few SNP.

Signals due to relationships are weak in a population with a very big Ne, and QTN profiles are narrow. Then the only choice for high accuracy is the identification of QTN or very close markers to QTN. Such an identification would require a very large SNP chip so that the individual SNP would fall within narrow QTN profiles or sequence information.

For populations with small effective population size, signals due to relationships would be strong, QTN profiles would be wide, the number of QTN with large effect would be small except in simulation studies or for unselected traits, and the identification of actual QTN would be hard. With small dataset, SNP selection may increase the accuracy of predictions due to the reduction in the dimensionality of the genomic information even inf the selected SNP are mostly due to signals due to relationships (Karaman *et al*. 2016; Lourenco *et al*. 2017; Pocrnic *et al*. 2019). With very large phenotypic data when signals due to noise would be small, large accuracy can be obtained without QTN identification since QTN can be accounted via QTN profiles. With medium datasets, the accuracy with SNP selection would be somewhat higher if the largest QTN or their markers can be identified; identification of actual SNP would require a sequence data.

A study by Fragomeni *et al*. (2019) provides a glimpse into factors affecting accuracy with sequence information. Reliabilities (squares of accuracy) were calculated for stature in Holsteins, where the genomic information included 54k generic SNP and 17k putative QTN on 27k bulls. Additionally, phenotypic information was available on 3M cows. Initial analyzes by GBLUP used pseudo-observations on the bulls. The base reliability with unselected 54k SNP was 69%, increased to 70% after including 17k putative SNP, and increased again to 71% with weighting the genomic relationship matrix; weighting is a form of SNP selection. After correcting the model for a different amount of information per bull, the reliabilities increased by 2%, with no advantage for weighting. After changing the model to ssGBLUP, where the phenotypes of cows were modeled directly, the reliability increased by 3%, again with no advantage for weighting. The study illustrates the point that an improved chip may improve the accuracy (1% in this case), SNP selection or weighting may compensate for an inferior model, and better modeling with more data has a much higher impact.

One of the goals of the 1000 bull genomes project was finding QTN based on sequence data, acknowledging that SNP in the regular chips (i.e., from 10k to 777k) are insufficient to capture the information about QTN (Hayes and Daetwyler 2019). Therefore, the central hypothesis behind discovering and using QTN in genomic evaluations is to help maximize prediction accuracies. However, the reported gains from sequence variants are only marginal (e.g., Veerkamp *et al*. 2016; Ros-Freixedes *et al*. 2022; Jang *et al*., 2022b). Summarizing earlier developments, small gains are likely for several reasons: the inability to identify the true causative QTN due to the wide QTN profiles in animal populations, few large QTN existing in selected populations, and GBLUP increasingly accounting for QTN with larger data.

## CONCLUSIONS

Manhattan plots are composed of signals from QTN, LD to QTN called QTN profile, relationships, and noise. The QTN profile is similar in shape to a pairwise linkage disequilibrium curve and has a width inversely proportional to the effective population size. With large effective population size, QTN profiles are narrow, relationships are weaker, and QTN identification is relatively easy with large phenotypic data. With a small effective population, signals due to QTN profiles are wide and confounded with strong signals due to relationships, resulting in limited resolution of GWAS and poor discovery rate. Genetic prediction in populations with large effective population size require high density SNP and identification of QTN or markers close to QTN. Genetic prediction in populations with small effective population size are sufficiently accurate with medium density SNP, and with large data account for QTN via QTN profiles, even without the actual QTN identification. QTN profiles provide a justification for showing Manhattan plots as a percentage of variance explained in moving windows. In such a case, the optimal window size for a population with N_e_=100 is 1-2 Mb wide.

## ACKNOWLEDGEMENT

The authors acknowledge Chris Gaynor (The Roslin Institute) for advice with AlphaSimR, Miguel Perez-Enciso (IRTA) and Martin Johnsson (Swedish University of Agricultural Sciences) for assistance with an earlier version of this study, and proofreading by Mary Kate Hollifield (The University of Georgia).

## Appendix

## Pairwise linkage disequilibrium curve and Stam segments

The expectation of pairwise linkage disequilibrium (r^2^) as a function of effective population size and distance from QTL (in Morgans), represented by c, was quantified by Sved (1971) as:

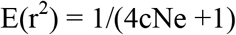

Assuming one Stam segment as 1/(4Ne), the curve can be rewritten as

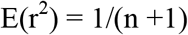

where n is the distance from the QTN in Stam segments. Thus, r^2^ declines to 0.7 for an interval of 1 Stam segment and to 0.2 for an interval of 8 Stam segments.

Assume q SNP per Stam segment, with the QTN represented by SNP 0. Assuming that the SNP value due to a single QTN is proportional to PLD,

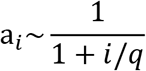

where q is a constant, and the variance assuming equal gene frequency is 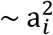. Then, the fraction of a QTN variance accounted for within t Stam segments is

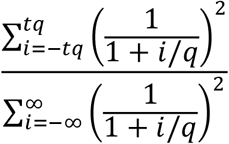

Numerical computations show that the interval of 2 segments explains about 50% of the QTN variance, 4 segments explain 66%, and 8 segments explain 80%.

